# Structural basis of ClC-3 inhibition by TMEM9 and PI(3,5)P_2_

**DOI:** 10.1101/2025.02.28.640562

**Authors:** Marina Schrecker, Yeeun Son, Rosa Planells-Cases, Sumanta Kar, Viktoriia Vorobeva, Uwe Schulte, Bernd Fakler, Thomas J. Jentsch, Richard K. Hite

## Abstract

The trafficking and activity of endosomes relies on the exchange of chloride ions and protons by members of the CLC family of chloride channels and transporters, whose mutations are associated with numerous diseases. Despite their critical roles, the mechanisms by which CLC transporters are regulated are poorly understood. Here, we show that two related accessory β-subunits, TMEM9 and TMEM9B, directly interact with ClC-3, -4 and -5. Cryo-EM structures reveal that TMEM9 inhibits ClC-3 by sealing the cytosolic entrance to the Cl^-^ ion pathway. Unexpectedly, we find that PI(3,5)P_2_ stabilizes the interaction between TMEM9 and ClC-3 and is required for proper regulation of ClC-3 by TMEM9. Collectively, our findings reveal that TMEM9 and PI(3,5)P_2_ collaborate to regulate endosomal ion homeostasis by modulating the activity of ClC-3.

## Main

The acidic organelles of the endolysosomal system require precise maintenance of their luminal ion concentrations, including protons, Cl^-^, and Ca^2+^, to ensure proper endosomal trafficking and function ^1-4^. Consequently, dysregulation of endolysosomal ion homeostasis is associated with numerous pathologies, ranging from kidney stones to neurodegeneration ^5-9^. Five members of the CLC family of Cl^-^ channels and Cl^-^/H^+^ exchangers, ClC-3, -4, -5, -6, and -7, play critical roles in endosomes and lysosomes, catalyzing the exchange of two Cl^-^ for one H^+ *2*,*10*^. CLC transporters are functional dimers with each protomer possessing an independent pathway for Cl^-^ transport ^2,11,12^. Located within the Cl^-^ ion pathway is a conserved glutamate residue, known as the gating glutamate, that is essential for the coupled transport of Cl^-^ and H^+ 11,13-16^. Structures of prokaryotic and eukaryotic CLC transporters have revealed that the gating glutamate can adopt several conformations that correspond to distinct states in the transport cycle ^2,11,12,14,17^. However, despite numerous structural investigations and their prominent roles in endolysosomal ion homeostasis, the mechanisms by which CLCs are regulated are poorly understood.

Many transport proteins are regulated by accessory β-subunits that can participate in membrane trafficking and/or directly modulate transport. CLCs have three known β-subunits: OSTM1, an obligatory β-subunit for the lysosomal ClC-7 transporter, barttin, an obligatory β-subunit for the plasma membrane channels ClC-Ka and ClC-Kb, and GlialCAM, a facilitative β-subunit for ClC-2 ^18-26^. How OSTM1 and barttin regulate the activities of the CLCs remains incompletely understood. Whether other β-subunits contribute to the trafficking and regulation of other CLCs is also unknown.

We recently found that the ClC-3, -4, and -5 clade of the endolysosomal CLC transporters require the accessory β-subunit TMEM9, which we will refer to as T9A, or its closely related homolog TMEM9B (T9B) for proper activity in cells and animals ^27^. Consistent with a recent report that T9B suppresses plasma membrane currents of ClC-3 and ClC-4 ^28^, we found that T9A and T9B strongly reduce the plasma membrane expression of members of the ClC-3 to ClC-5 clade ^27^. Moreover, we found that T9A and T9B directly inhibit CLC ion transport through a mechanism that requires their C-terminal domains ^27^. To resolve the mechanism by which T9A and T9B regulate the activity of ClC-3, -4, and -5, we determined cryo-EM structures of human ClC-3 alone and in complex with T9A, finding that the C-terminal domain acts a reversible pore blocker and that the regulation of ClC-3 by T9A requires the endolysosomal signaling lipid PI(3,5)P_2_.

## Results

### Structure of human ClC-3

To determine how T9A and T9B regulate ClC-3, -4, and -5, we collected cryo-EM images of human ClC-3 alone and in complex with human T9A in the absence of exogenous ligands. We focused on ClC-3 because it forms stable complexes with T9A and T9B, whereas detergent-solubilized ClC-4 and ClC-5 only partially associate with T9A and T9B (**Extended Data Fig. 1**). Analysis of the images of ClC-3 yielded a C2 symmetric reconstruction of dimeric ClC-3 at a resolution of 2.5 Å (**Fig. 1a-b, Extended Data Fig. 2 and Extended Data Table 1**). Each ClC-3 protomer is comprised of a cytosolic N-terminal domain (NTD), a transmembrane domain (TMD), and a cytosolic domain (CD) (**Fig. 1c**). The NTD contains helix A and an extended loop that is sandwiched between the TMD and the CD. The TMD of ClC-3 adopts the canonical CLC fold with distinct a Cl^-^ ion pathway passing through the center of each protomer. However, in contrast to the continuous helical conformation of helix B in bovine ClC-K, human ClC-2, human ClC-6, or human ClC-7, helix B of ClC-3 is separated into two segments by a 32-residue insertion that we will refer to as the helix B insertion ^25,29-31^ (**Extended Data Fig. 3**). The ClC-3 helix B insertion is stabilized by a pair of disulfide bonds and adopts partially ordered conformation. The ClC-3 CD contains two tandem CBS domains that establish an ATP-binding site, in which no density corresponding to a bound adenine nucleotide was observed, indicating that we captured a ligand-free structure ^32,33^ (**Fig. 1c**).

**Fig. 1:**
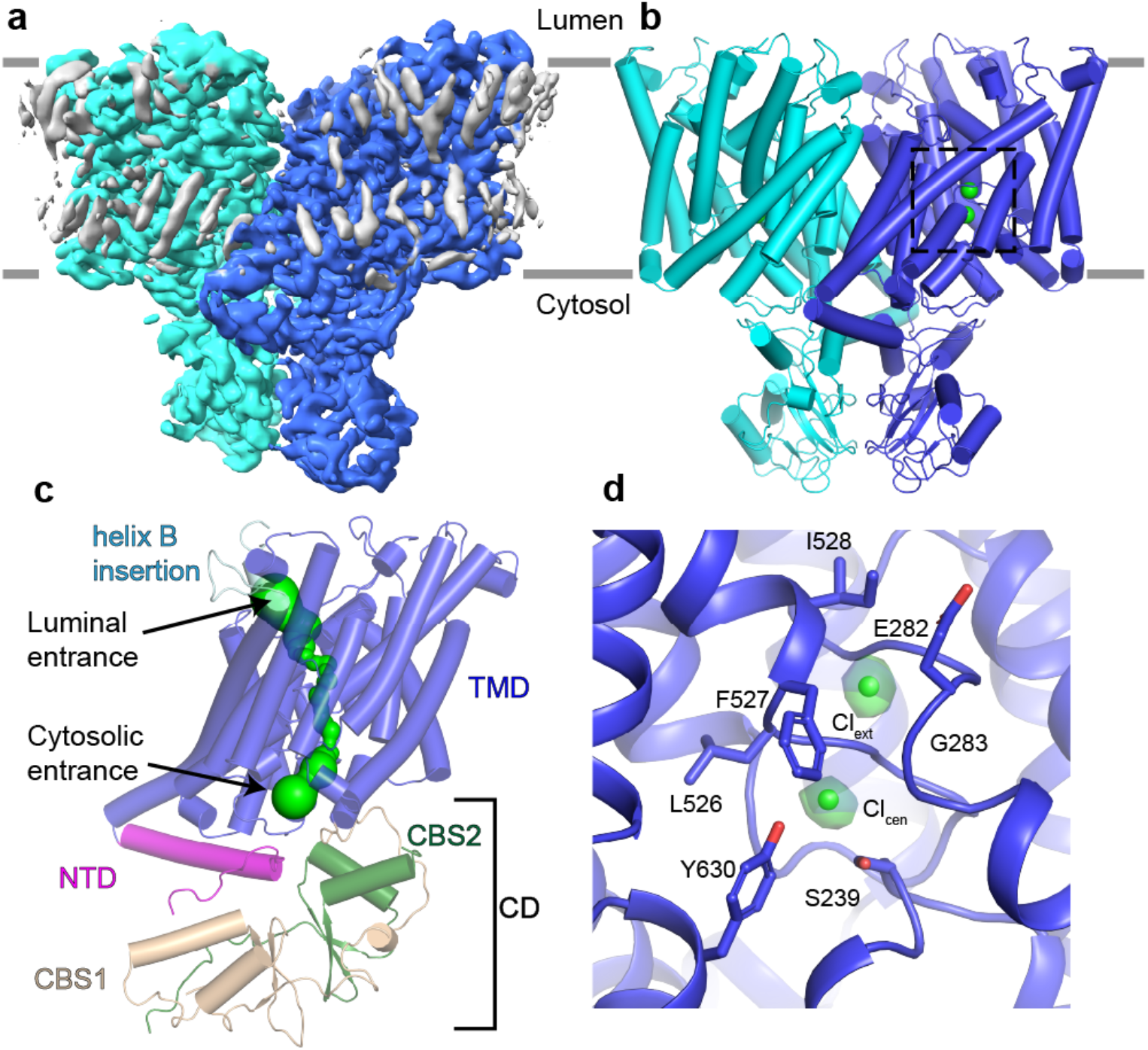
Structure of human ClC-3. (**a-b**) Cryo-EM density map (A) and fitted atomic model (B) of ClC-3 colored by protomer. Bound Cl^-^ ions are shown as green spheres. Dashed box in B corresponds to panel D. **(c)** Domain architecture of ClC-3 with the NTD colored in magenta, the TMD colored in blue, the helix B insertion colored in light blue, CBS-1 colored in tan and CBS-2 colored in dark green. The Cl^-^ ion pathway is shown as a green surface. **(d)** Cl^-^-binding sites in the Cl^-^ ion pathway of one protomer. Densities corresponding to Cl^-^ ions are shown as green isosurfaces and contoured at 4.0 *σ*.

The entrances at both the luminal and cytosolic sides of the Cl^-^ ion pathway are solvent-accessible. Density peaks occupy the central and external Cl^-^-binding sites that we assigned as bound Cl^-^ ions (**Fig. 1d**). The central Cl^-^-binding site is formed by the side chains of ^239^Ser and ^630^Tyr and the backbone nitrogen of ^526^Leu, and the external binding site is formed by the backbone nitrogen atoms of ^282^Glu, ^283^Gly, ^527^Phe and ^528^Ile. Non-protein densities are also present near the internal Cl^-^-binding site, which were modeled as ordered water molecules as their identities were unclear. The conserved gating glutamate, ^282^Glu, adopts an outward conformation, establishing a constriction with a minimum radius of 0.4 Å between the external Cl^-^-binding site and the solvated luminal entrance to the Cl^-^ ion pathway. Overall, these features are consistent with reported structures of other CLC transporters, including a recently reported structure of mouse ClC-3 ^33^, indicating that this structure of human ClC-3 reflects a transport-competent state.

### Structure of ClC-3 in complex with T9A

Analysis of images of ClC-3 in complex with T9A revealed several distinct classes (**Fig. 2a and Extended Data Fig. 2 and 4**). In all classes, ClC-3 adopts a dimeric arrangement that closely resembles the conformation adopted by ClC-3 alone, whereas the densities corresponding to T9A differed (**Extended Data Fig 4**). We identified a symmetric class in which the luminal, transmembrane, and cytosolic domains of both T9A protomers are resolved, which we will refer to as ClC-3/T9A, and a symmetric class in which no densities corresponding to T9A are resolved, which we will refer to as ClC-3/noT9A (**Fig. 2a and Extended Data Fig. 4**). We also identified asymmetric classes in which the TMD of T9A is resolved but the cytosolic and/or luminal domains are disordered (**Extended Data Fig. 4**). Due to its completeness, we will focus our discussion primarily on ClC-3/T9A, which achieved a resolution of 2.9 Å with C2 symmetry imposed (**Extended Data Table 1**).

**Fig. 2:**
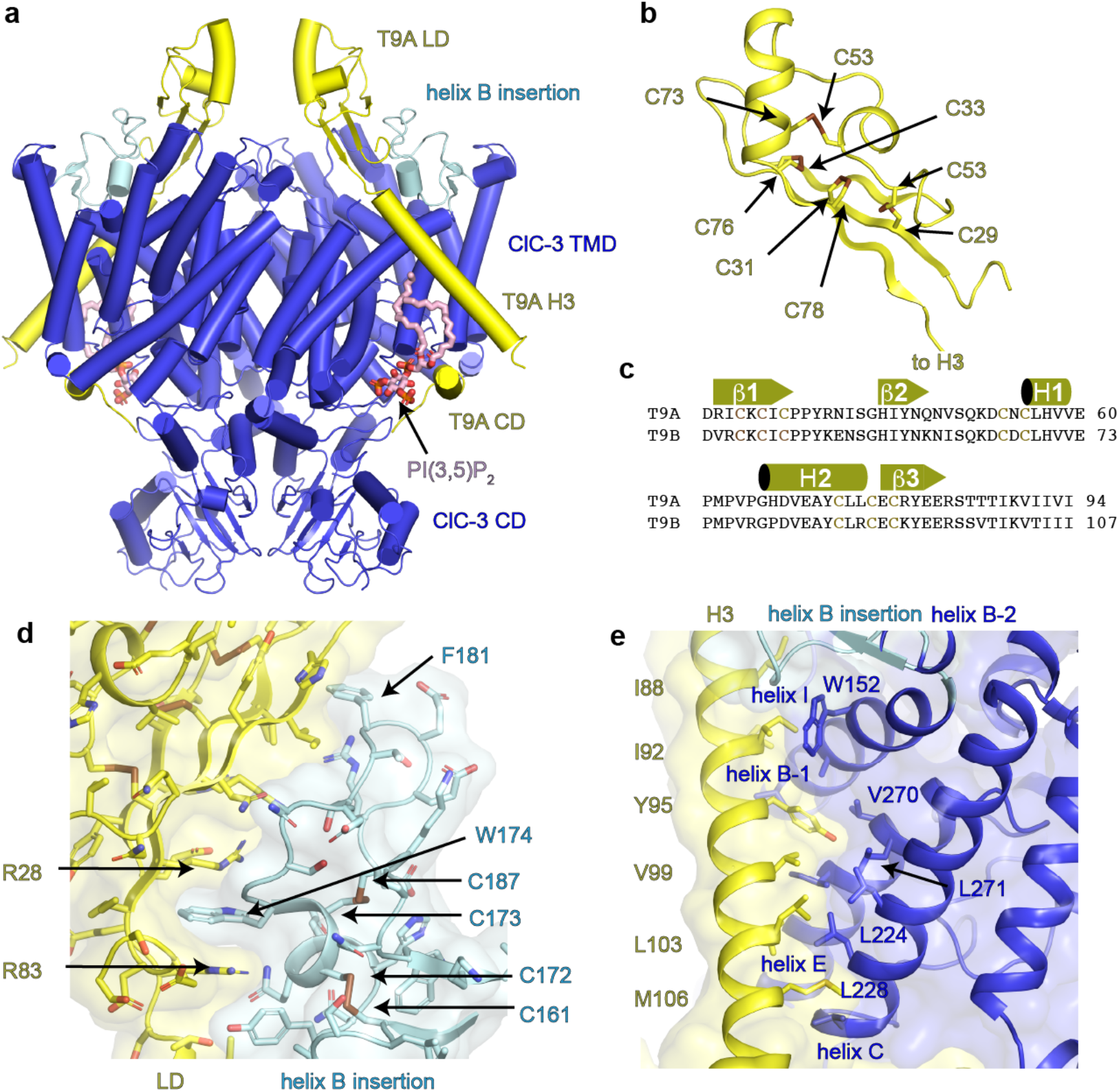
Structure of human ClC-3 in complex with T9A. **(a)** ClC-3/T9A structure of ClC-3 colored by subunit with T9A in yellow, ClC-3 in blue and the ClC-3 helix B insertion in light blue. PI(3,5)P_2_ is shown in pink. **(b)** Structure of T9A LD with residues that form disulfide bonds shown as sticks. **(c)** Sequence alignment of human T9A and T9B LDs. Secondary structural elements and residues that form disulfide bonds are highlighted. **(d)** Interaction between T9A LD and the helix B insertion of ClC-3. **(e)** Interaction between H3 of T9A and the TMD of ClC-3

In ClC-3/T9A, T9A wraps almost entirely around ClC-3, interacting with the luminal, transmembrane, and cytosolic faces of ClC-3. Notably, we do not observe any interactions between the two T9A protomers. The T9A luminal domain (LD) is comprised of a three-stranded β-sheet and two short *α*-helices, a fold that is stabilized by four disulfide bonds (**Fig. 2b**). The residues that establish these disulfide bonds are conserved in human T9B, suggesting that the fold of the LD is conserved (**Fig. 2c**). The T9A LD interacts primarily with the ClC-3 helix B insertion forming both polar and non-polar interactions, including the embedding of ^181^Phe of ClC-3 into a hydrophobic groove on T9A and an extended *π*-stacking interaction between ^28^Arg and ^83^Arg of T9A with ^174^Trp of ClC-3 (**Fig. 2a,d**). The interactions with T9A stabilize the ClC-3 helix B insertion in a more ordered state in ClC-3/T9A, allowing us to model the entire domain. The sequence of the helix B insertion is highly conserved in ClC-3, ClC-4 and ClC-5 but not in ClC-6 or ClC-7, suggesting that the helix B insertion is a characteristic feature among the CLC transporters that interact with T9A and T9B (**Extended Data Fig. 3**).

The single membrane-spanning helix of T9A, helix H3, extends along the periphery of the ClC-3 TMD, forming interactions with helices B, C, E, and I of ClC-3 (**Fig. 2e**). Despite being only a single helix, its interaction with ClC-3 buries a large surface area. The interactions between helix H3 and the TMD of ClC-3 are almost exclusively hydrophobic, with residues such as ^92^Ile and ^95^Tyr of T9A occupying hydrophobic grooves on the surface of ClC-3.

The T9A CD consists of the H4 helix that extends along the cytosolic face of the ClC-3 TMD and four residues at the extreme C-terminus, which we define as the C-terminal domain (CTD) (**Fig. 2a and 3a**). The T9A CD is connected to the H3 helix via an extended linker that is disordered in the structure and contains a stretch of negatively charged residues and several phosphorylation sites that contribute to the trafficking of ClC-3 and T9A ^27^. The H4 helix interacts with the cytosolic ends of helices C, D, E, J, and R, the D-E linker, and the J-K linker. Among the residues on the T9A H4 helix that interact with ClC-3 are ^164^Trp, which contacts ^232^Phe and ^260^Trp of ClC-3, and Phe176, which is inserted into a groove between ^230^Lys, ^234^Pro, ^427^Pro and ^521^Lys of ClC-3 (**Fig. 3b**). Additionally, the side chain of ^172^Arg of T9A interacts with the backbone oxygen atoms of ^230^Lys and ^231^Val of ClC-3. Collectively, these interactions position the H4 helix such that the four-residue CTD can extend from helix H4 along the side of ClC-3 towards the cytosolic domain, where ^183^Ser of T9A interacts with the backbone of ^88^Phe and the sidechain of ^798^Lys of ClC-3 (**Fig. 3c**).

**Fig. 3:**
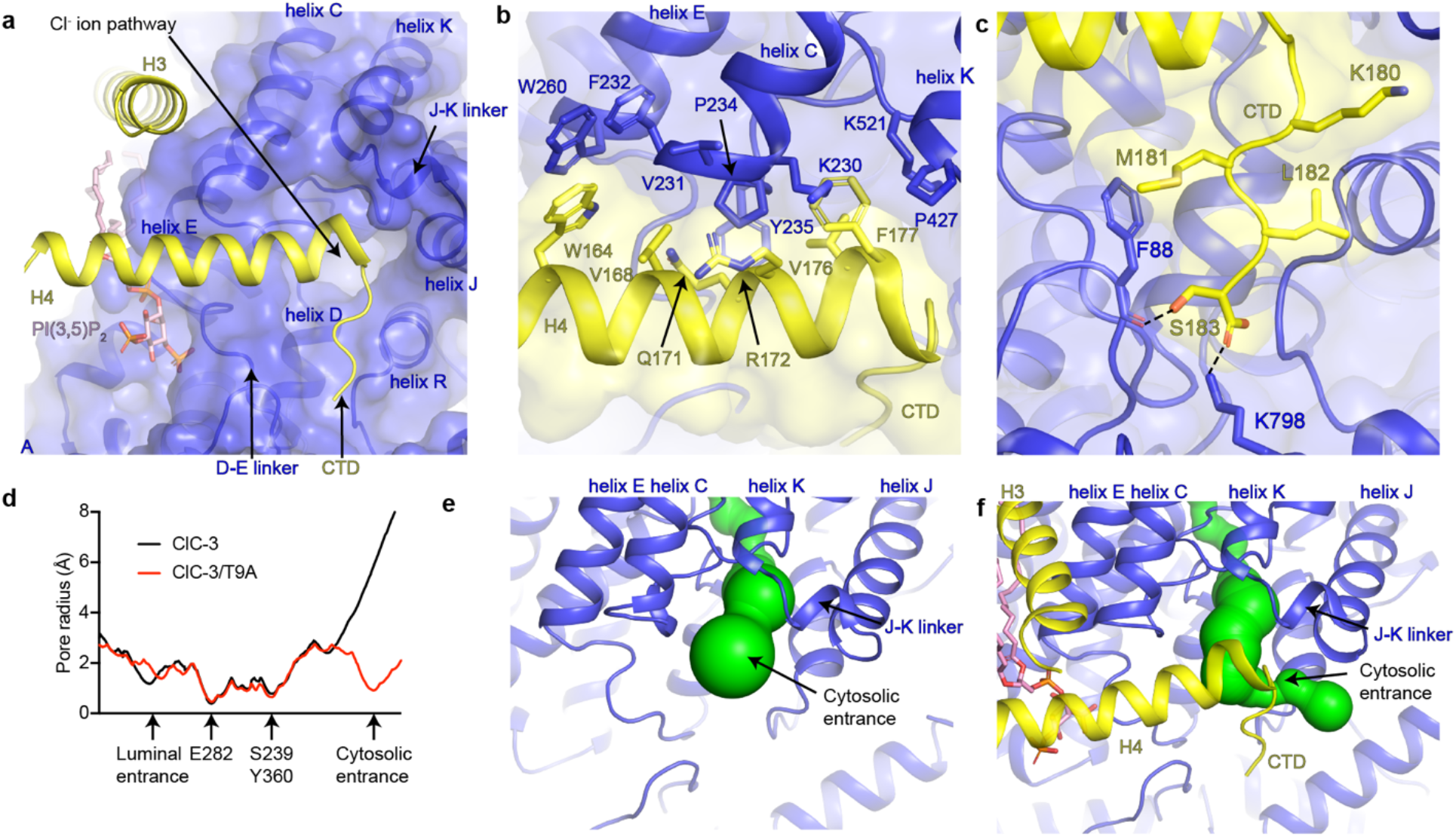
The cytosolic domain of T9A plugs the Cl^-^ pathway. **(a)** The T9A CD, consisting of helix H4 and the four-residue CTD, binds to ClC-3 near the cytosolic entrance to the Cl^-^ ion pathway. **(b)** Interactions between helix H4 of T9A and ClC-3. **(c)** Interactions between the four-residue CTD of T9A and ClC-3. **(d)** Pore radius plot of Cl^-^ ion pathways of ClC-3 (black) and ClC-3/T9A (red). (**e-f**) The cytosolic entrance of the Cl^-^ ion pathway is open in ClC-3 (e) and sealed in ClC-3/T9A (f). Cl^-^ ion pathways are shown as green surfaces.

### T9A and OSTM1 bind to distinct interfaces

To further investigate the specificity of T9A and T9B for ClC-3, -4, and -5 and of OSTM1 for ClC-7, we compared ClC-3/T9A with a structure of ClC-7/OSTM1 ^25^. Although T9A and OSTM1 are both single-pass transmembrane proteins with disulfide-stabilized LDs that serve as obligatory β-subunits, T9A and OSTM1 bind to distinct surfaces on their respective CLCs (**Extended Data Fig. 5**) ^22,27^. For example, whereas the T9A LD binds to the ClC-3 helix B insertion, the LDs of OSTM1 form few interactions with ClC-7 and instead form an extensive homodimeric interface that helps to stabilize CLC-7/OSTM1 in the harsh environment of the lysosome ^25^. The TMDs of T9A and OSTM1 also bind in dissimilar manners: T9A traverses along the side of the ClC-3 TMD in a highly tilted manner, while OSTM1 binds in upright manner to the extreme periphery of the ClC-7 TMD. Lastly, in contrast to the extensive interactions formed between the T9A CD and ClC-3, no evidence for an interaction between the OSTM1 CD and ClC-7 has been observed in the reported structures ^25,26^. Thus, despite their common origin, ClC-3, -4, and -5 have evolved unique features that enable interactions with T9A and T9B compared to those that have evolved in ClC-7 to facilitate its interactions with OSTM1.

### The T9A CD plugs the cytosolic entrance to the pore of ClC-3

To understand how T9A inhibits ClC-3, we compared the structure of ClC-3 alone with the ClC-3/T9A structure. As mentioned previously, T9A does not alter the global conformation of ClC-3 in the ClC-3/T9A structure. T9A also had no effect on the conformation of the gating glutamate, ^282^Glu, or the positions of the ions in the central and external Cl^-^-binding sites in ClC-3/T9A (**Extended Data Fig. 6**). Rather, T9A inhibits ClC-3 by sealing the cytosolic entrance to the Cl^-^ pathway. Whereas both entrances of the Cl^-^ ion pathway are solvent accessible in the absence of T9A, the H4 helix and CTD of T9A constrict the cytosolic entrance to a minimum radius of 0.9 Å, which is too narrow to accommodate a Cl^-^ ion (**Fig. 3d-f**). We therefore propose that the T9A CD prevents Cl^-^ ions from accessing the cytosolic entrance to the Cl^-^ ion pathway in a manner that directly analogous to a pore blocker of an ion channel and we will use this terminology. Consistent with this proposal, mutations in either T9A or ClC-3 that disrupt the interface between the T9A CD and ClC-3 diminish the ability of T9A to inhibit ClC-3 ^27^. Mutations in the T9B CD similarly diminished its ability to inhibit ClC-3, indicating that the T9B CD also acts as pore blocker ^27^. Likewise, mutations in the CD of T9A and T9B suppress the inhibitory effects of T9A and T9B on ClC-4 and ClC-5 ^27^.

In addition to ClC-3/T9A, in which the CDs of both protomers bind ClC-3, we also resolved structures in which the CDs of one or both T9A protomers are disordered (**Extended Data Fig. 4**). Despite T9A being present in these classes, dissociation of the T9A CD results in an opening of the cytosolic entrance to the Cl^-^-permeation pathway, suggesting that the CD functions as a reversible pore blocker and must be bound to ClC-3 to inhibit transport (**Extended Data Fig. 6**).

### PI(3,5)P_2_ is required for the inhibition of ClC-3 by T9A

In ClC-3/T9A, we observed a non-protein density near the interface between ClC-3 and the T9A helix H4. We modelled this density as a co-purified phosphatidylinositol (3,5) bis-phosphate (PI(3,5)P_2_), a lipid synthesized from PI(3)P by PIKfyve in endosomes and lysosomes ^34^ (**Figs. 3a and 4a**). Densities corresponding to PI(3,5)P_2_ are also resolved in the other reconstructions of ClC-3 determined in the presence of T9A, including ClC-3/noT9A (**Extended Data Fig. 4**). Although the densities are less well resolved in ClC-3/noT9A than in ClC-3/T9A, potentially suggesting lower occupancy, the PI(3,5)P_2_ density is entirely absent in the ClC-3 only reconstruction and in the recent reconstructions of mouse ClC-3 ^33^, suggesting that T9A contributes to PI(3,5)P_2_ binding.

In ClC-3/T9A, PI(3,5)P_2_ interacts with both ClC-3 and the T9A CD. The phosphate groups at the 1 and 3 positions of the PI(3,5)P_2_ inositol ring bind to ClC-3, whereas the phosphate at the 5 position interacts with the T9A CD. The phosphate at the 1 position is coordinated by the backbone nitrogen atoms of ^259^Lys and ^260^Trp of ClC-3, and the side chain of ^259^Lys of ClC-3. The phosphate at the 3 position binds to the backbone nitrogen of ^255^Gly of ClC-3 and the side chains of ^254^Arg, ^293^Asn, ^298^Tyr and ^310^Lys of ClC-3. The 5 position phosphate interacts with the side chain of ^163^Arg from the T9A H4 helix. Thus, PI(3,5)P_2_ appears to serve as a bridge between ClC-3 and the T9A CD, raising the possibility that PI(3,5)P_2_ contributes to the regulation of ClC-3.

We next investigated the role of PI(3,5)P_2_ in the T9A-mediated inhibition of ClC-3. ClC-3 activity in endosomes can be visualized by monitoring the enlargement of endosomal vacuoles marked by Venus-ClC-3 ^27,35,36^. Venus-ClC-3 expression in HeLa cells led to the appearance of enlarged endosomal vacuoles, a phenotype that was suppressed by co-transfection with T9A, consistent with the inhibitory effect of T9A on ClC-3 ^27^ (**Fig. 4b-c**). To ensure specificity for ClC-3, all experiments were performed in cell lines deficient for ClC-7, whose inhibition by PI(3,5)P_2_ blocks lysosomal vacuolization ^37^. Treating ClC-7-deficient HeLa cells with apilimod, which inhibits PI(3,5)P_2_ generation by PIKfyve ^34^, suppressed the inhibitory effect of T9A, restoring endosomal enlargement in wild-type ClC-3-expressing cells (**Fig. 4d**). Notably, apilimod had no impact on endosomal vacuolization in cells expressing the transport deficient (*td*) E339A mutant of ClC-3 (**Fig. 4e**). Taken together, these results demonstrate that PI(3,5)P_2_ formation is required for the inhibitory effects of T9A on ClC-3 activity in endosomes.

**Fig. 4:**
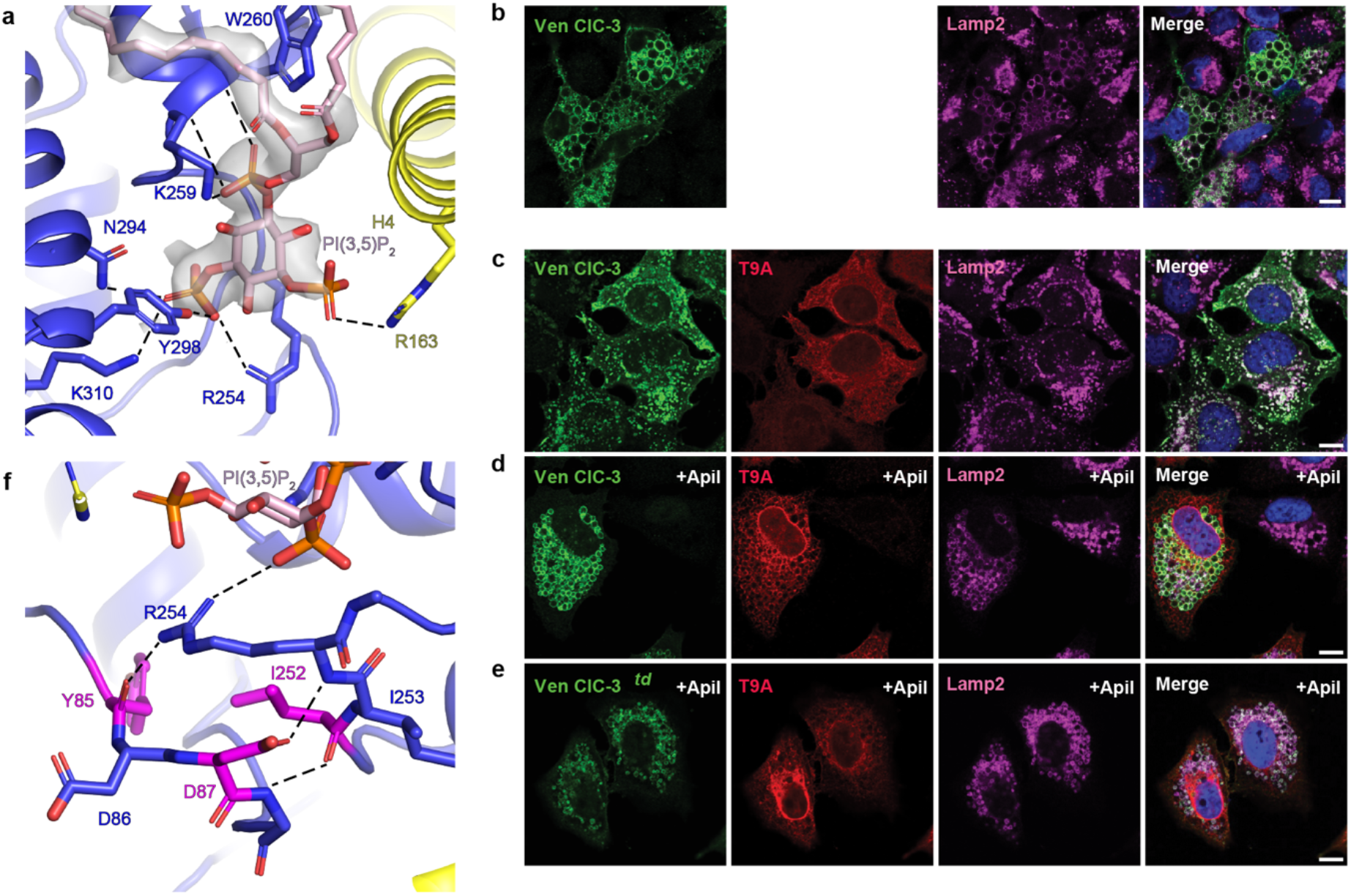
Inhibition of ClC-3 by T9A requires PI(3,5)P_2_. (**a**) PI(3,5)P_2_ binding site at the interface between ClC-3 and helix H4 of T9A. Dashed lines correspond to hydrogen bonding interactions. Density corresponding to PI(3,5)P_2_ is displayed as a grey isosurface and contoured at 2.5 *σ*. (**b-e**) ClC-3-dependent endosomal vacuole generation is inhibited by apilimod. Confocal imaging of Venus-ClC-3a (green), T9A (red) and Lamp2 (magenta) from ClC-7 KO HeLa cells expressing Venus-ClC-3a (B), expressing Venus-ClC-3a and T9A (C), expressing Venus-ClC-3a and T9A treated with 100 nM apilimod for 4 hours (D) and expressing Venus-ClC-3a^td^ and T9A treated with 100 nM apilimod for 4 hours (E). Merged image at right. (**f**) Network of interactions that stabilize the interface between ClC-3 and PI(3,5)_2_. Residues whose mutation are associated with human diseases are highlighted in magenta.

Numerous mutations associated with human disease have been identified in *CLCN3*, which encodes ClC-3, and *CLCN4*, which encodes ClC-4 ^6,38^. Although some of the mutations affect ion transport by ClC-3 in isolation ^6,38^, other mutations modulate the inhibitory effects of T9A and T9B on ClC-3 and can only be observed when they are co-expressed ^27^. To gain insights into how disease-associated mutations in *CLCN3* and *CLCN4* perturb the inhibitory effects of T9A and T9B, we mapped the mutations onto ClC-3/T9A. Although many of the mutations occurred in residues near the interface with T9A, several were located near the PI(3,5)P_2_ binding site, highlighting the importance of PI(3,5)P_2_ in the regulation of CLCs (**Fig. 4f and Extended Data Fig. 7**). For example, the backbone oxygen atom of ^85^Tyr of ClC-3, which is mutated to cysteine in neurological disease ^6^, interacts with the side chain of ^254^Arg of ClC-3 that binds the PI(3,5)P_2_ head group. ^87^Asp, which corresponds to ^29^Asp in ClC-4 that is mutated to glutamate in a neurological disease ^38^, and ^252^Ile in ClC-3, which is mutated to threonine in a neuropathy patient ^6^, also interact with ^254^Arg. Collectively, our results indicate that T9A and PI(3,5)P_2_ are dynamic regulators of endosomal CLC function and that disruption of the T9A-CLC interface can lead to disease ^27^.

## Discussion

In this study, we demonstrate that ClC-3, and likely ClC-4 and ClC-5, are regulated by T9A and PI(3,5)P_2_. The mode of regulation enables a mechanistic understanding of pathological mutations in the genes encoding CLCs that can lead to disease. In contrast to mutations that disrupt ion transport or alter the conformational landscape of ClC-3, we show that mutation of residues mediating interactions with accessory β-subunits can also lead to dysregulated CLC activity ^27^.

The CDs of T9A and T9B inhibit ClC-3 by reversibly blocking the cytosolic entrance to the Cl^-^ ion pathway to inhibit Cl^-^/H^+^ exchange. Recently, a structure of the related Cl^-^ channel, ClC-2, was resolved in which an inhibitory peptide from its own N-terminus was found to block the cytosolic entrance of the Cl^-^ ion pathway (**Extended Data Fig. 8**) ^30,39,40^. Thus, in a manner analogous to K^+^ channels, for which numerous peptides have evolved to bind to block the extracellular entrance to the K^+^-permeation ion pathway, our work establishes that peptides from distinct origins can regulate CLCs by occluding the cytosolic entrance to the Cl^-^ ion pathway. Future work may identify additional examples of protein interaction partners that can regulate the activity of CLCs by binding to the cytosolic entrance to the Cl^-^ ion pathway.

Unexpectedly, we found that inhibition of ClC-3 requires the endolysosomal signaling lipid, PI(3,5)P_2_, which stabilizes the T9A CD in the inhibitory bound conformation. PI(3,5)P_2_ has also been shown to inhibit ClC-7 ^37^, although the mechanism of inhibition is unknown and the PI(3,5)P_2_ binding site has not been definitively identified. Suggestively, the PI(3,5)P_2_ binding site in ClC-3 partially overlaps with a site near the cytosolic end of helix E in ClC-7 that binds the related phosphatidylinositol, PI(3)P ^25^ (**Extended Data Fig. 9**). Future studies will be necessary to determine if the overlap of the binding sites is coincidental or if PI(3,5)P_2_ inhibits ClC-7 through a similar mechanism.

Phosphoinositol lipids, such as PI(3,5)P_2_, regulate the activity of numerous transport proteins including ClC-7, TRPML1, and TPC1/2 ^37,41,42^. Structures of TRPML1 and TPC1 in the presence and absence of PI(3,5)P_2_ reveal that PI(3,5)P_2_ binding induces global conformational changes that alter the ion permeation pathways ^43,44^. Here, we establish a new paradigm where PI(3,5)P_2_ binding gates ion permeation by acting as a molecular glue to stabilize the interaction between a transport protein and an accessory β-subunit. This mode of regulation would enable rapid and reversible control of CLC activity in endosomes through the generation and subsequent breakdown of PI(3,5)P_2_ by the kinase PIKfyve and the PtdIns(3,5)P2 5-phosphatase, FIG4, respectively ^45,46^. Through the ability of PI(3,5)P_2_ to regulate diverse ion channels and transporters in endosomes and lysosomes, PIKfyve and FIG4 serve as master regulators of endolysosomal ion homeostasis, directly contributing to the regulation of H^+^, Cl^-^, Na^+^, and Ca^2+^.

## Methods

### Fluorescence-detection size-exclusion chromatography (FSEC)

Genes encoding human ClC-3 (P51790), human ClC-4 (P51793), human ClC-5 (P51795), human ClC-6 (P51797), and human ClC-7 (P51798) were synthesized by Twist Biosciences and subcloned into BacMam expression vectors with N-terminal mCerulean tags fused with a linker containing a PreScission protease site ^47^. Genes encoding human T9A (Q9P0T7), human T9B (Q9NQ34), and human OSTM1 (Q86WC4) were synthesized by Twist Biosciences and subcloned into BacMam expression vectors with a C-terminal mVenus tag fused with a linker containing a PreScission protease site ^47^. All construct sequences were validated by Sanger sequencing.

800 μl of Expi293F cells with a cell density of 3×10^6^ were dispensed into a 96 deep-well plate (Corning, P-DW-20-C-S). 0.8 ug DNA and 2.4 μg of PEI 25K (Polysciences, Inc) were each mixed with 50 μl of Opti-MEM Reduced Serum Medium (Gibco) and incubated for 5 minutes at room temperature. For complex transfection, equal amounts of the plasmid encoding relevant N-terminal mCerulean-tagged CLC and the plasmid encoding C-terminal mVenus-tagged T9A or T9B were used. After incubation, DNA was combined with PEI 25K (Polysciences, Inc) and incubated for 20 min at room temperature and then used for transfection. After 24 hr incubation at 37°C with continuous agitation, valproic acid sodium salt (Sigma-Aldrich, P4543) was added to a final concentration of 2.2 mM, and cells were allowed to grow at 37°C for an additional 24 hr before harvesting. Cell pellets were washed in phosphate-buffered saline solution (PBS) and flash frozen in liquid nitrogen. Expressed proteins were solubilized in 200 µl buffer containing 2% lauryl maltose neopentyl glycol (LMNG) (Anatrace, NG310), 20 mM HEPES pH 7.5, 150 mM KCl supplemented with protease-inhibitor cocktail (1 mM PMSF, 2.5 mg/mL aprotinin, 2.5 mg/mL leupeptin, 1 mg/mL pepstatin A) and DNaseI. Solubilized proteins were separated by centrifugation at 21,130 g for 45 min. Separated proteins were injected to and monitored by fluorescence-detection size-exclusion chromatography on a Superose 6 Increase 10/300 GL (GE healthcare) in a buffer composed of 0.02% glyco-diosgenin (GDN) (Anatrace, GDN101), 150 mM KCl, 20 mM HEPES pH 7.5, and 1 mM dithiothreitol (DTT) (Goldbio, DTT10). mCerulean fluorescence was monitored at 433/475 nm for excitation/emission wavelength, respectively.

### Protein expression and purification

For expressing ClC-3 in complex T9A, the gene encoding human T9A (Q9P0T7) was synthesized by Twist Biosciences and subcloned into BacMam expression vector ^47^, of which a stop codon was introduced after the T9A sequence to ensure that T9A was not fused to a fluorophore with a linker containing a PreScission protease site. The N-terminal mCerulean tagged ClC-3 construct described above was used for ClC-3/T9A complex expression. Equal amounts of the plasmid encoding N-terminal mCerulean-tagged ClC-3 and the plasmid encoding T9A without a fused fluorophore were mixed 1:3 (w/w) with PEI 25 K (Polysciences, Inc) for 20 min and then used to transfect Expi293 cells. Once the cell density reached 3 x 10^6^ cells/ml, 1 mg plasmid and 3 mg PEI 25 K were used to transfect a 1L cell culture. For ClC-3 expression by itself, the plasmid encoding C-terminal mCerulean-tagged ClC-3 was used to transfect HEK293S GnTi– cells (ATCC: CRL-3022) at cell desnity 2 x 10^6^ cells/ml using the same protocol described above.

After 24 hr incubation at 37°C, valproic acid sodium salt (Sigma-Aldrich, P4543) was added to a final concentration of 2.2 mM, and cells were allowed to grow at 37 °C for an additional 24 hr before harvesting. Cell pellets were washed in phosphate-buffered saline solution and flash frozen in liquid nitrogen. Membrane proteins were solubilized in 2% LMNG (Anatrace, NG310), 0.2% Cholesteryl Hemisuccinate Tris Salt (CHS) (Anatrace, CH210), 20 mM HEPES (pH 7.5), 150 mM KCl supplemented with protease-inhibitor cocktail (1 mM PMSF, 2.5 mg/mL aprotinin, 2.5 mg/mL leupeptin, 1 mg/mL pepstatin A), and DNase I. Solubilized proteins were separated by centrifugation 135,557 g for 40 min at 4°C, followed by binding to anti-GFP nanobody resin for 1.5 hr at 4°C, which had previously been equilibrated with Size Exclusion Chromatography (SEC) buffer containing 0.02% glyco-diosgenin (GDN, Anatrace), 50 mM Tris-HCl (pH 8), 150 mM KCl, and 2 mM DTT. Anti-GFP nanobody affinity chromatography was performed by 2-3 column volumes of washing with SEC buffer followed by overnight PreScission digestion and elution with the SEC buffer. The eluted protein sample was concentrated to a volume of 250 *μ*l using CORNING SPIN-X concentrators (100 kDa cutoff) (Corning, 431491), followed by centrifugation 21,130 g for 15 min. Concentrated protein was further purified by size exclusion chromatography on Superose 6 Increase 10/300 GL (GE healthcare) in SEC buffer. Peak fractions were pooled and concentrated using CORNING SPIN-X concentrators (100 kDa cutoff) to a concentration specified in the electron microscopy sample preparation method section below.

### Electron microscopy sample preparation and data acquisition

For ClC-3, 3 μl of 3mg/ml purified protein in the absence of any exogenous ligands was applied to glow-discharged Au 400 mesh QUANTIFOIL R1.2/1.3 holey carbon grids (Quantifoil). The grids were plunged into liquid nitrogen-cooled liquid ethane using an FEI Vitrobot Mark IV (FEI Thermo Fisher). The freezing process occurred at 4°C under 100% humidity, with blotting times ranging from 2 to 3.5 s and a waiting time of 10 s. The grids were transferred to a 300 keV FEI Titan Krios microscope equipped with a K3 summit direct electron detector (Gatan). Images were captured with SerialEM ^48^ in super-resolution mode at a magnification of 29,000x, corresponding to a pixel size of 0.413 Å. The dose rate during imaging was set at 15 electrons/pixel/s, and the defocus range was from -0.7 to -2 µm. The image acquisition duration was 3 s, consisting of 0.05 subframes (total of 60 subframes), resulting in a cumulative dose of 66 electrons/Å^2^.

For the ClC-3 in complex with T9A complex, 3 μl of 5 mg/ml purified protein in the absence of any exogenous ligands was applied to glow-discharged Au 400 mesh QUANTIFOIL R1.2/1.3 holey carbon grids (Quantifoil) and then plunged into liquid nitrogen-cooled liquid ethane with an FEI Vitrobot Mark IV (FEI Thermo Fisher). The sample was frozen at 4°C with 100% humidity, using blotting times of 2.5 s, blotting force of 2, and a waiting time of 10 s. Grids were transferred to a 300 keV Titan Krios microscopy equipped with a Falcon 4i direct detector and a SeletrisX energy filter (Krios G4) (Thermo scientific). The images were recorded with EPU software in pixel size of 0.725 Å. Dose rate was 11.51 electrons/pixel/s, and the defocus range was -0.5 to -1.5 μm. Images were recorded for 2.93 s with frame time of 0.003256 s (total 900 frames), corresponding to a total dose of 59.63 electrons/Å^2^. The nominal magnification was 165,000X. Energy filter slit width was 10 eV.

### Electron microscopy data processing

#### ClC-3

4,672 of 60-frame super-resolution movies (0.413 Å/pixel) of ClC-3 collected from on grid were gain corrected, Fourier cropped by two (0.826 Å/pixel) and aligned using whole-frame and local motion correction algorithms by cryoSPARC v3.2.0 ^49^. Blob-based autopicking in cryoSPARC was implemented to select initial particles, resulting in stacks of 1,446,105 particles. False-positive selections and contaminants were removed by iterative rounds of heterogenous classification using the initial 3D reconstruction generated using the ab initio reconstruction in cryoSPARC as well as several decoy classes generated from noise particles via ab initio reconstruction in cryoSPARC v4.2.1 ^49^. After particle polishing in Relion 3.1.2 ^50^, a non-uniform refinement of the resulting particles yielded a reconstruction of 507,952 particles at 2.94Å. Iterative rounds of heterogenous classification followed by a C2-symmetry non-uniform refinement in cryoSPARC v4.2.1 with local CTF estimation and higher order aberration correction resulted in a final reconstruction of 219,662 particles at 2.54Å.

#### ClC-3 in complex with T9A

19,481 of nine-hundred-frame movies (0.725 Å/pixel) of ClC-3 in complex with T9A were collected from two grids using a Falcon 4i direct detector with a SelectrisX energy filter at a slit width of 10 eV (ThermoFisher Scientific). The movies were fractionated to 40 and aligned using whole-frame and local motion correction algorithms in cryoSPARC v4.2.1 ^49^. Blob-based autopicking in cryoSPARC was implemented to select initial particles, resulting in stacks of 5,874,584 and 4,262,456 particles. A subset of particles with best 2D averages was selected after performing 2D classification in cryoSPARC. False-positive selections and contaminants of the subset particles were removed by iterative rounds of heterogenous classification using the final human ClC-3 reconstruction as well as several decoy classes generated from noise particles via ab initio reconstruction in cryoSPARC v4.2.1 ^49^. After performing 2D classification of the resulting particle stacks, particles with the best 2D averages were then selected to train topaz picking in cryoSPARC v4.2.1 ^51^, resulting in stacks of 1,316,391 and 1,065,143 particles. False-positive selections and contaminants of the topaz-picked particles were excluded through iterative rounds of heterogenous classification using the same inputs as above, resulting in stacks of 331,753 and 51,335 particles. False-positive selections and contaminants of the initial blob-based auto-picked particles were also removed via iterative rounds of heterogenous classification using the same inputs, resulting in stacks of 408,736 and 342,430 particles. Duplicate particles from the blob-based auto-picked particles and topaz-picked particles from the same micrograph dataset were removed. After performing reference-based motion correction of the duplicate-removed particles, the particles from different micrograph dataset were combined. The combined particles were subjected to C2 symmetry non-uniform refinement in cryoSPARC v4.2.1 with per-particle defocus and global CTF estimations, resulting in a reconstruction of 1,057,848 particles at 2.71Å. The particle stack was then classified via 3D classification in cryoSPARC v4.2.1 with C1 symmetry, using a focus mask that covers the density of T9A. A distinct class of ClC-3 dimer with no T9A density was identified. A C2-symmetry non-uniform refinement of the class resulted in reconstruction of 148,161 particles at 2.90 Å. Particles of the remaining classes were further classified by iterative rounds of 3D classification with the T9A focus mask. Three classes with distinct T9A densities were identified. A C1-symmetry non-uniform refinement of the first class, of which the CTD density of T9A was absent in both T9A and the LD density was absent in one of the two T9A, resulted in reconstruction of 91,755 particles at 3.01 Å. A C1-symmetry non-uniform refinement of the second class, of which LD and CTD densities of both T9A were present, resulted in reconstruction of 94,011 particles at 2.86 Å. A C1-symmetry non-uniform refinement of the third class, of which LD and CTD densities of T9A was present only in one of the two T9A, resulted in reconstruction of 71,754 particles at 3.16 Å. Other classes consisted of mixed populations of poorly classified particles.

### Model building and coordinate refinement

#### ClC-3

The final reconstruction was subjected to density modification using the two unfiltered half-maps with a soft mask in Phenix ^52^, yielding an improved density map at 2.45 Å. Human ClC-3 was manually built into the density modified map in COOT v0.9.6 ^53^, followed by iterative rounds of real space refinement in phenix v1.21.1-5286 ^54^and manual rebuilding in COOT v0.9.6 ^53^.

#### ClC-3 in complex with T9A

The final reconstructions were subjected to density modification using the two unfiltered half-maps with a soft mask in Phenix ^52^. The structure of human ClC-3 alone was docked into the maps and manually rebuilt in COOT v0.9.6 to fit the density ^53^. Human T9A was then manually built into the density modified map. The models were refined by iterative rounds of real space refinement in phenix v1.21.1-5286 ^54^and manual rebuilding in COOT v.0.9.6 ^53^.

#### ClC-3 vacuolization assay

3× 10^4^ Hela *CLCN7* KO cells ^7^ seeded on 10 mm glass coverslips (1.5H) (Langenbrinck GmbH) were co-transfected with expression plasmids of the short transcription ClC-3 variant (ClC-3a) fused N-terminally to Venus fluorescent protein (Venus-ClC-3a) plus T9A (*28*). As controls, Venus-ClC-3a was substituted by a transport deficient mutant (E281A, according to short transcript), *td*), or T9A was substituted by CD4. 48 hours later, cells were treated for 4 h with 100 nM apilimod (Cat. SML2974, Sigma-Aldrich) or DMSO as vehicle control in complete medium at 37 °C and 5% CO_2_. Cells were then washed twice in cold PBS and fixed with methanol for 15 min. Cells were extensively washed twice with PBS, blocked and permeabilized in 3% sterile-filtered goat serum (NGS) (Cat. P30-1002, PAN-Biotech), 2% BSA and 0.1% saponin at room temperature for 1 h. Cells were then incubated overnight at 4 °C with chicken anti-GFP antibody (1: 500) (Cat# GFP-1020, Aves Lab), guinea pig anti-T9A (T9AC2) (1:1000) (Pineda Antibody Service, Berlin) and mouse anti-Lamp-2 (H4B4) (Cat. Ab25631, Abcam) primary antibodies solution in 3% BSA, 0.05% saponin). Next, cells were extensively washed in 0.05% saponin-PBS, incubated with secondary antibodies coupled to different fluorophores and 1 μg/ml DAPI at room temperature for 1 h, then washed and mounted on slides using Fluoromount-G (SouthernBiotech) and allowed to dry. Images were acquired using a LSM880 Zeiss confocal microscope using a 63X NA 1.4 oil-immersion lens.

#### Statistics and Reproducibility

All FSEC experiments and were independently performed three times, with the results being similar. All ClC-3a vacuolization experiments were independently performed two times, with the results being similar. Cryo-EM images were collected from 1-2 grids per sample condition. The number of collected images for each condition is indicated in Extended Data Figure 2.

## Figures

Figures were prepared with PyMol 2.5.3. (www.pymol.org), ChimeraX 1.5 ^55^, GraphPad Prism 9 (www.graphpad.com), Clustal Omega 1.2.3. ^56^, and MOLE ^57^.

## Data Availability

Cryo-EM maps have been deposited in the EMDB under accession codes EMD-47070 [http://www.ebi.ac.uk/emdb/EMD-47070] (ClC-3), EMD-47066 [http://www.ebi.ac.uk/emdb/EMD-47066] (ClC-3/noT9A), EMD-47067 [http://www.ebi.ac.uk/emdb/EMD-46067] (ClC-3/T9A, T9A Protomer A and B: Complete), EMD-47068 [http://www.ebi.ac.uk/emdb/EMD-47068] (ClC-3/T9A, T9A Protomer A: No CD, T9A Protomer B: No LD, No CD) and EMD-47069 [http://www.ebi.ac.uk/emdb/EMD-47069] (ClC-3/T9A, T9A Protomer A: Complete, T9A Protomer B: No LD, No CD). Atomic coordinates have been deposited in the PDB under accession codes 9DO0 [https://doi.org/10.2210/pdb9DO0/pdb] (ClC-3), 9DNW [https://doi.org/10.2210/pdb9DNW/pdb] (ClC-3/no T9A), 9DNX [https://doi.org/10.2210/pdb9DNX/pdb] (ClC-3/T9A, T9A Protomer A and B: Complete), 9DNY [https://doi.org/10.2210/pdb9DNY/pdb] (ClC-3/T9A, T9A Protomer A: No CD, T9A Protomer B: No LD, No CD) and 9DNZ [https://doi.org/10.2210/pdb9DNZ/pdb] (ClC-3/T9A, T9A Protomer A: Complete, T9A Protomer B: No LD, No CD). The atomic coordinates of previously published structures of bovine ClC-K [https://doi.org/10.2210/pdb5TQQ/pdb], human ClC-2 [https://doi.org/10.2210/pdb8TA4/pdb], human ClC-6 [https://doi.org/10.2210/pdb8JPJ/pdb], and human ClC-7/OSTM1 complex [https://doi.org/10.2210/7JM7/pdb] were used in this study.

## Acknowledgments

We thank M. J. de la Cruz for help with data acquisition; the Memorial Sloan Kettering Cancer Center HPC group for assistance with data processing and the members of the laboratories for comments on the manuscript. This study is supported in the Hite lab in part by NIGMS R01-GM141553 and NIH-National Cancer Institute Cancer Center Support Grant P30-CA008748. This study is supported in the Jentsch lab in part by the European Research Council (ERC) Advanced Grant 740537 (VolSignal) and the Deutsche Forschungsgemeinschaft (DFG) JE164/12-2 and under Germany’s Excellence Strategy–EXC-2049 Project ID 390688087 (Neurocure). This study is supported in the Fakler lab in part by grants of the Deutsche Forschungsgemeinschaft (DFG; FA 332/15-1, 16-1 and 21-1 and the CRC/TRRs 1453 and 152). M.S. is supported by the Walter Benjamin Programme of the DFG.

## Author contributions

Conceptualization, M.S., Y.S., T.J. and R.H.; Methodology M.S., Y.S., R.P-C., V.V., S.K., U.S., B.F., T.J. and R.H., Formal Analysis, M.S., Y.S., R.P-C., V.V., S.K., U.S., B.F., T.J. and R.H.; Investigation, M.S., Y.S., R.P-C., V.V., S.K., U.S., B.F., T.J. and R.H.; Writing – Original Draft, M.S., Y.S. and R.H.; Funding Acquisition, M.S., B.F., T.J. and R.H.

