## Supplementary data for "Structural basis of ClC-3 inhibition by TMEM9 and PI(3,5)P_2_"

**Extended Data:**

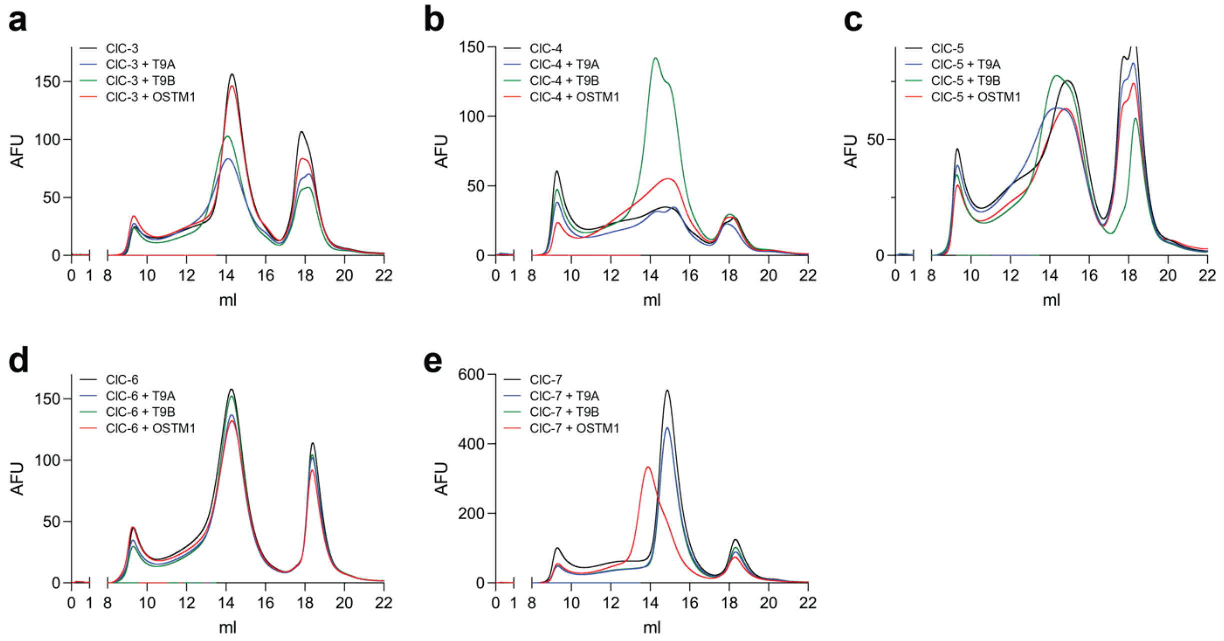

**Extended Data Figure 1: T9A and T9B form complexes with ClC-3, ClC-4 and ClC-5.** Expi293F cells were transfected with different DNA constructs. After protein purification, the fluorescence of mCerulean-tagged protein was monitored at an excitation/emission wavelength of 433/475 nm, respectively.

(**a**) FSEC profiles of mCerulean-ClC-3 (black), mCerulean-ClC-3 co-expressed with mVenus-T9A (blue), mCerulean-ClC-3 co-expressed with mVenus-T9B (green), and mCerulean-ClC-3 co-expressed with mVenus-OSTM1 (red).

(**b**) FSEC profiles of mCerulean-ClC-4 (black), mCerulean-ClC-4 co-expressed with mVenus-T9A (blue), mCerulean-ClC-4 co-expressed with mVenus-T9B (green), and mCerulean-ClC-4 co-expressed with mVenus-OSTM1 (red).

(**c**) FSEC profiles of mCerulean-ClC-5 (black), mCerulean-ClC-5 co-expressed with mVenus-T9A (blue), mCerulean-ClC-5 co-expressed with mVenus-T9B (green), and mCerulean-ClC-5 co-expressed with mVenus-OSTM1 (red).

(**d**) FSEC profiles of mCerulean-ClC-6 (black), mCerulean-ClC-6 co-expressed with mVenus-T9A (blue), mCerulean-ClC-6 co-expressed with mVenus-T9B (green), and mCerulean-ClC-6 co-expressed with mVenus-OSTM1 (red).

(**e**) FSEC profiles of mCerulean-ClC-7 (black), mCerulean-ClC-7 co-expressed with mVenus-T9A (blue), mCerulean-ClC-3 co-expressed with mVenus-T9B (green), and mCerulean-ClC-7 co-expressed with mVenus-OSTM1 (red).

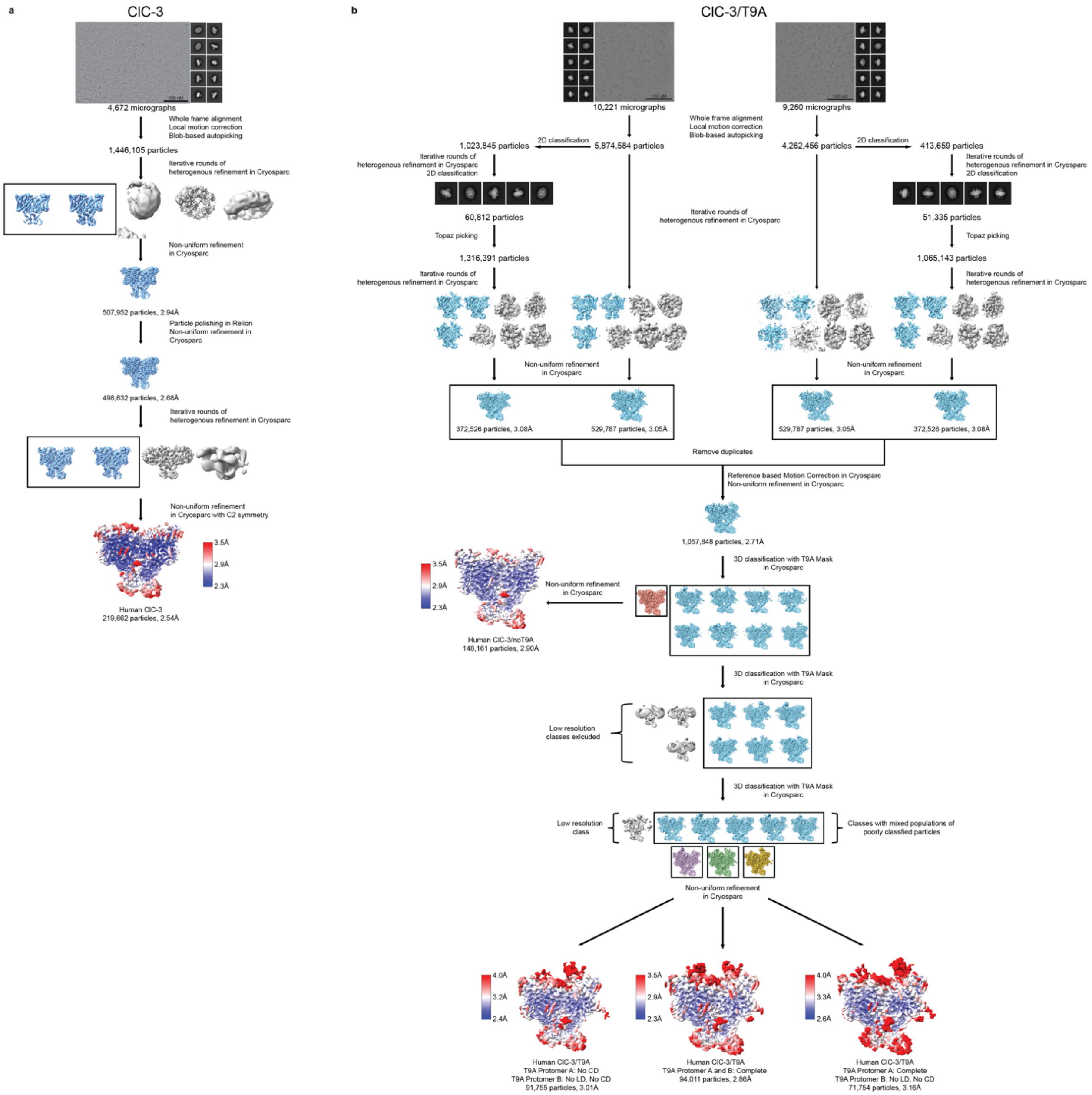

**Extended Data Fig. 2: Cryo-EM analyses of ClC-3 (a) and ClC-3 in complex with T9A (b).**

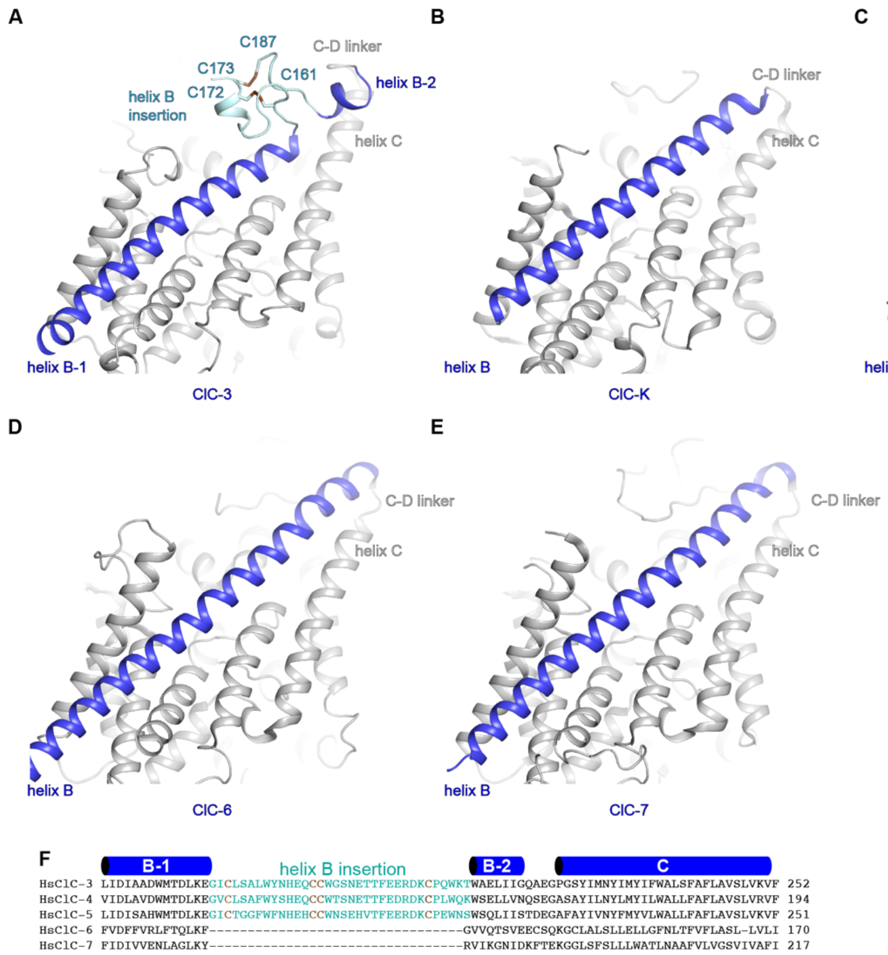

**Extended Data Fig. 3: Conformation of helix B in mammalian CLC structures.**
(**a-e**) Cartoon depiction of TMD of human ClC-3 (a), bovine ClC-K (b, PDB 5TQQ), human ClC-2 (c, PDB 8TA4), human ClC-6 (d, PDB 8JPJ) and human ClC-7 (e, PDB 7JM7). Helix B is colored in blue, the helix B insertion in ClC-3 is colored in light blue and all other residues are colored in grey. Residues in ClC-3 that form disulfide bonds are shown as sticks.
(**f**) Sequence alignment of helix B of human ClC-3, ClC-4, ClC-5, ClC-6, and ClC-7. Secondary structural elements and residues that form disulfide bonds in ClC-3 are highlighted.

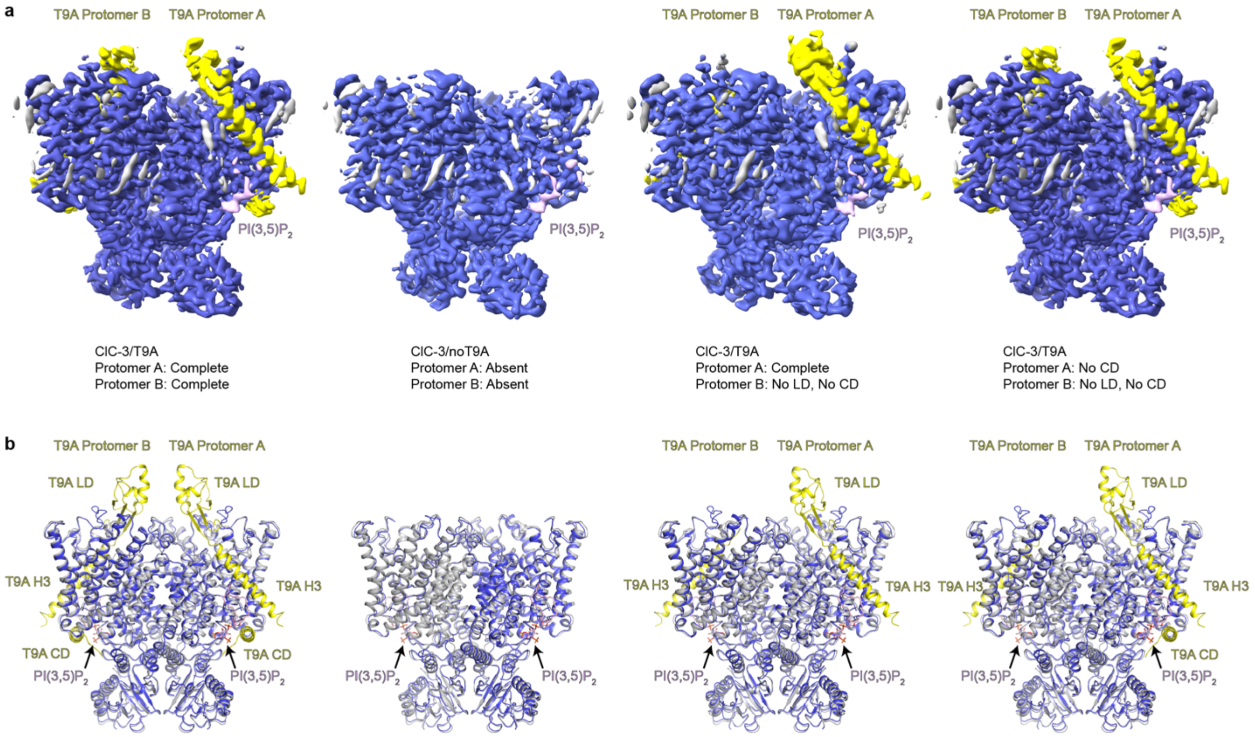

**Extended Data Fig. 4: Structures of ClC-3 in the presence of T9A.**
(**a**) Cryo-EM maps of ClC-3/T9A, ClC-3/noT9A and two classes of ClC-3 in complex with T9A in which the T9A protomers are only partially ordered, colored by subunit. Densities corresponding to ClC-3 are colored blue, T9A are colored yellow and PI(3,5)$P_2$ are colored pink.
(**b**) Superposition of ClC-3 alone (grey) with ClC-3/T9A, ClC-3/noT9A and two classes of ClC-3 in complex with T9A in which the T9A protomers are only partially ordered (colored by subunit).

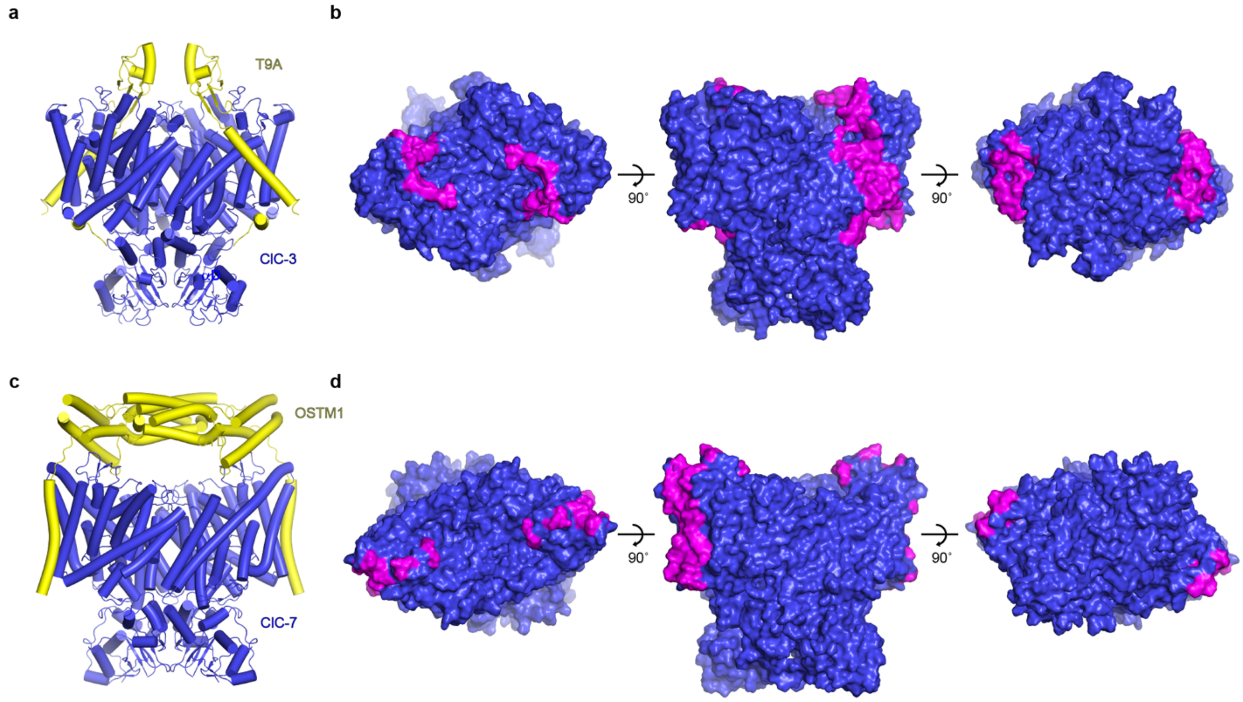

**Extended Data Fig. 5: Comparison of ClC-3/T9A and ClC-7/OSTM1 complexes.**
(**a,c**) Structures of ClC-3/T9A (a) and ClC-7/OSTM1 (c, PDB: 7JM7), colored by subunit.
(**b,d**) Surface representation of ClC-3 (b) and ClC-7 (d), viewed from the cytosol (left), from within the plane of the membrane (middle) and lumen (right). Residues that interact with T9A or OSTM1 are colored in magenta.

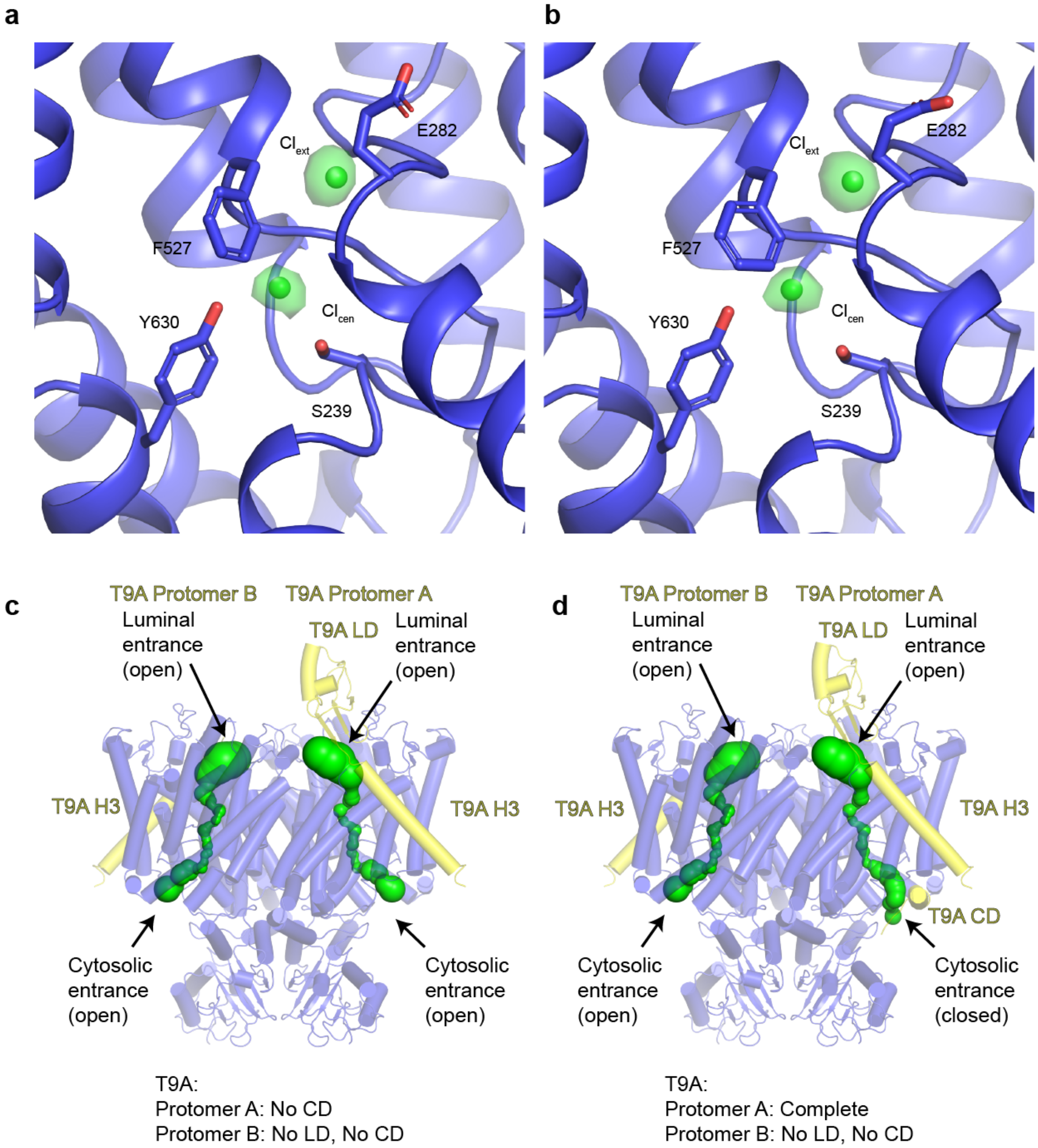

**Extended Data Fig. 6: Cl$^-$ ion pathway of ClC-3 in the presence of T9A.**
(**a-b**) Cl$^-$-binding sites in the Cl$^-$ ion pathway of one protomer of ClC-3/T9A (a) or ClC-3/noT9A (b). Densities are shown as green isosurfaces and contoured at 4.0 s.
(**c-d**) Cl$^-$ ion pathways of two classes of ClC-3 in complex with T9A in which the T9A protomers are only partially ordered, depicted as green surfaces.

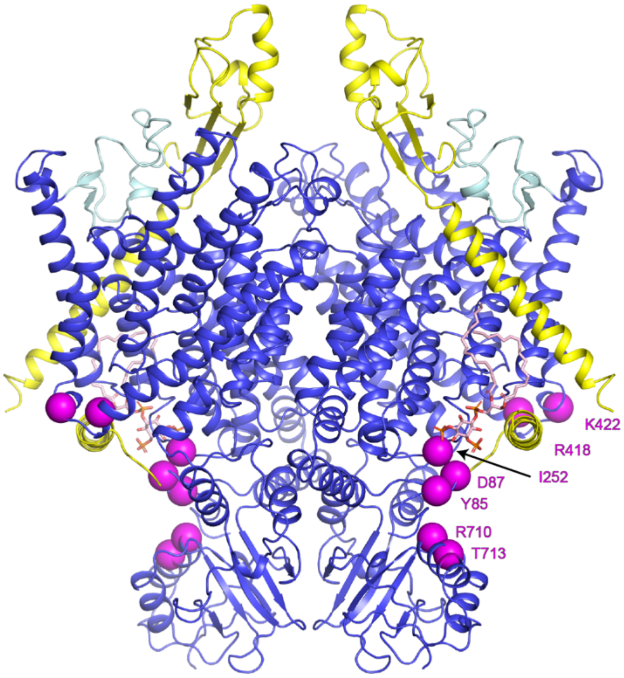

**Extended Data Fig. 7: Disease-associated mutations in ClC-3/T9A.** Residues whose mutation are associated with disease and weaken transport inhibition by T9A [28] are depicted as magenta spheres.

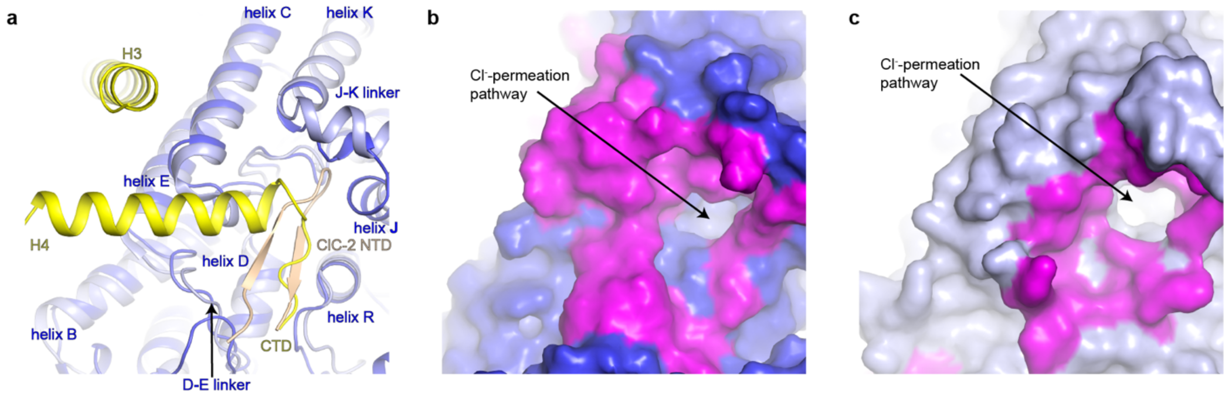

**Extended Data Fig. 8: Comparison of ClC-3/T9A and ClC-2.**

(**a**) Superposition of the region surrounding of the cytosolic entrances to the $Cl^-$ ion pathways of ClC-3/T9A and ClC-2 (PDB 8TA4). ClC-3 is colored blue, T9A is colored yellow, ClC-2 is colored light blue, and the ClC-2 NTD is colored sand.

(**b-c**) Surface representation of the region surrounding the cytosolic entrances to the $Cl^-$ ion pathways of CLC3 (b) and ClC-2 (c), with residues that interact with T9A or the N-terminus of ClC-2 colored in magenta.

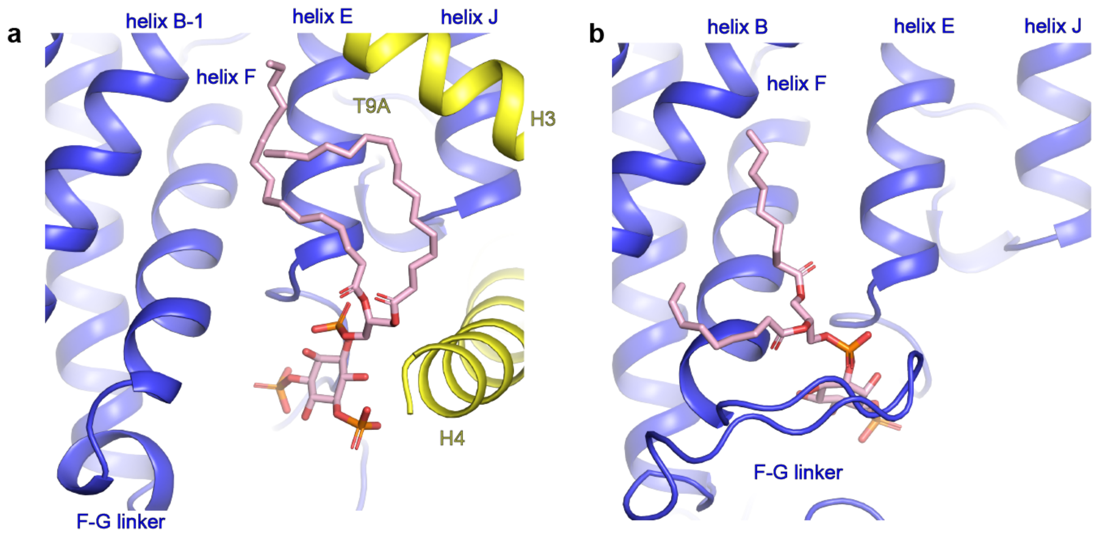

**Extended Data Fig. 9: Phosphatidylinositol binding sites in ClC-3/T9A and ClC-7/OSTM1.**
(**a**) PI(3,5)$P_2$ binding site in ClC-3/T9A.
(**b**) PI(3)P binding site in ClC-7/OSTM1 (PDB 7JM7).

| | #1 ClC-3 (EMD-47070) (PDB 9DO0) | #2 ClC-3/noT9A (EMD-47066) (PDB 9DNW) | #3 ClC-3/T9A<br><br>T9A Protomer A and B: Complete<br><br>(EMD-47067) (PDB 9DNX) | #4 ClC-3/T9A<br><br>T9A Protomer A: No CD<br>T9A Protomer B: No LD, No CD<br><br>(EMD-47068) (PDB 9DNY) | #5 ClC-3/T9A<br><br>T9A Protomer A: Complete<br>T9A Protomer B: No LD, No CD<br><br>(EMD-47069) (PDB 9DNZ) |
|---|---|---|---|---|---|
| **Data collection and processing** | | | | | |
| Detector | Gatan K3 | TFS Falcon4i | TFS Falcon4i | TFS Falcon4i | TFS Falcon4i |
| Magnification | 29,000X | 165000X | 165000X | 165000X | 165000X |
| Voltage (kV) | 300 | 300 | 300 | 300 | 300 |
| Energy filter slit width (eV) | | 10 | 10 | 10 | 10 |
| Electron exposure (e–/Å$^2$) | 66 | 59.63 | 59.63 | 59.63 | 59.63 |
| Defocus range (μm) | -0.7 to -2 | -0.5 to -1.5 | -0.5 to -1.5 | -0.5 to -1.5 | -0.5 to -1.5 |
| Super-resolution pixel size (Å) | 0.413 | | | | |
| Final pixel size (Å) | 0.826 | 0.725 | 0.725 | 0.725 | 0.725 |
| Symmetry imposed | C2 | C2 | C2 | C1 | C1 |
| Initial particle images (no.) | 1,446,105 | 10,137,040 | 10,137,040 | 10,137,040 | 10,137,040 |
| Final particle images (no.) | 219,662 | 148,161 | 94,011 | 91,755 | 71,754 |
| Map resolution (Å)<br>FSC threshold | 2.54<br>0.143 | 2.90<br>0.143 | 2.86<br>0.143 | 3.01<br>0.143 | 3.16<br>0.143 |
| **Refinement** | | | | | |
| Model resolution (Å)<br>0.5 FSC threshold | 2.51 | 2.79 | 2.84 | 2.96 | 3.13 |
| Map sharpening *B* factor (Å$^2$) | -30 | -30 | -30 | -30 | -30 |
| Model composition | 11,130 | 11,208 | 13,280 | 12,326 | 12,524 |

| | | | | | |
|---|---|---|---|---|---|
| Non-hydrogen atoms<br>Protein residues<br>Ligands | 1,394<br>8 | 1,399<br>10 | 1,656<br>10 | 1,540<br>10 | 1,562<br>10 |
| Mean *B* factors ($Å^2$)<br>Protein<br>Ligand<br>Water | 58.0<br>52.1<br>26.4 | 26.1<br>15.5 | 67.4<br>52.3 | 46.0<br>35.2 | 63.8<br>53.3 |
| R.m.s. deviations<br>Bond lengths (Å)<br>Bond angles (°) | 0.002<br>0.426 | 0.002<br>0.449 | 0.003<br>0.477 | 0.002<br>0.476 | 0.002<br>0.499 |
| Validation<br>MolProbity score<br>Clashscore<br>Poor rotamers (%) | 1.07<br>2.83<br>0.09 | 1.24<br>4.54<br>1.03 | 1.41<br>5.35<br>1.43 | 1.40<br>5.96<br>1.24 | 1.35<br>5.34<br>1.22 |
| Ramachandran plot<br>Favored (%)<br>Allowed (%)<br>Disallowed (%) | 98.48<br>1.52<br>0.00 | 98.48<br>1.52<br>0.00 | 98.77<br>1.23<br>0.00 | 98.95<br>1.05<br>0.00 | 98.44<br>1.56<br>0.00 |

**Extended Data Table 1. Cryo-EM data collection and refinement statistics.**
